# Benchmarking Peptide Spectral Library Search

**DOI:** 10.64898/2026.01.08.698418

**Authors:** Hao Xu, Nuno Bandeira

## Abstract

Spectral library search (SLS) is a major approach for peptide identification from tandem mass spectrometry data, with performance depending substantially on the accuracy of the underlying Spectrum-Spectrum Matching (SSM) scoring functions. However, detailed comparative studies remain limited by the absence of comprehensive benchmark datasets. We propose new methods to build SSM scoring functions benchmarks and construct a benchmark dataset with (i) *eight* query spectrum sets with varying noise level for 476,063 precursors, and (ii) *three* spectral libraries with experimental, de-noised and predicted spectra for 3,065,819 precursors. We evaluate common spectrum preprocessing scenarios and SSM scoring functions, including SpectraST and EntropyScore. Results revealed this remains an important open problem, with the best recall for still assessed to be poor at just ∼70%, with SpectraST performing best for spectra with little-to-no noise, while JS-divergence showed superior noise resistance. Conversely, Cosine and Entropy score performed substantially worse, with Projected-Cosine performing especially poorly in most cases, with overall performance and relative ranking depending quite significantly on the minimum number of matching peaks. The benchmark dataset (MSV000095946/PXD056205) supports testing and development of new SSM scoring functions and the proposed benchmark construction approach provides an extensible foundation for additional types of SSM evaluation.

## INTRODUCTION

Proteomics is a wide-ranging field of study dedicated to the comprehensive identification and quantification of peptides and proteins in biological systems.^1,2^ Mass Spectrometry is a critical tool in this field, used to identify and quantify a sample’s components (i.e., peptides) by producing spectra of the masses and relative abundances of ionized peptides (MS1 spectra) or peptide fragments (tandem, MS2 or MS/MS spectra). Spectral library search^3^ (SLS) is a major approach for peptide identification from tandem mass spectrometry data, offering a complementary approach to conventional sequence database search.^4,5,6^ While database search methods identify peptides by matching query spectra against the peptide sequences from public sequence databases (based on their characteristic peptide fragment ion patterns,^5^ SLS methods identify peptides by comparing the query spectra against libraries of reference peptide spectra (usually assembled from previously identified spectra^7^) based on the similarity of peaks and intensity patterns in query and reference spectra.^8^

Previous studies have demonstrated the superior performance of SLS methods over database search methods.^9,10^ However, a significant limitation of SLS methods is their inability to detect novel peptides when reference spectra are absent in the library.^8^ Despite this, the emergence of spectrum prediction models^11,12^ has the potential to overcome this limitation, effectively transforming database search into SLS of predicted spectra. Recent studies^13,14^ have shown that utilizing predicted intensity information can enhance the accuracy when searching against a Homo sapiens UniProt sequence database in current database search methods, such as MS-GF+^5^ and Percolator.^15^ However, the performance of the Spectrum-Spectrum Match (SSM) scoring functions used by SLS tools on the predicted spectrum library was systematically evaluated.

Over the past decades, substantial research on SLS methods has been devoted to enhancing accuracy in matching tandem mass spectra.^3,16–23^ These methods typically include two critical steps in their peptide identification pipeline: spectrum preprocessing and spectrum-spectrum matching (SSM). Preprocessing methods are utilized to select spectrum peaks and transform peak intensities. Different preprocessing methods can be combined to enhance SLS search performance when preparing query and library spectra. In the SSM step, an SSM scoring function calculates a score that reflects the similarity between two spectra. These scores are then used to select the best match from potential candidates. The most widely used SSM functions in current SLS methods are based on the inner product of vector representations for matched peaks from two spectra (equivalent to cosine similarity when vectors have Euclidian norm 1), with variations in vector normalization. Recently, methods based on relative entropy / Jensen-Shannon divergence^24^ have demonstrated performance gains over inner-product-based methods in matching small molecule spectra,^23^ and have also shown promise in peptide identification.^25,26^ An SLS method may sometimes include a third step, in which SSM scores are adjusted using additional SSM information, such as the SSM scores for all candidate library matches for a particular query spectrum (used in SpectraST^21^). In addition, the analysis of chimeric/mixture spectra of co-eluting peptides is also usually based on Projected-Cosine, a version of the Cosine SSM scoring function that computes the cosine between a reference spectrum R and only those peaks in the query spectrum Q that match to a peak in R (essentially using cosine to assess whether R is contained in Q). Projected-cosine was initially designed for Data-Dependent Acquisition (DDA) spectra in MSPLIT^19^ and later extended for Data-Independent Acquisition (DIA) in MSPLIT-DIA.^27^ Even though Projected-Cosine is primarily designed and used for chimeric/mixture spectra, we also need to consider its performance in the simplest case of matching SSMs of single peptide spectra. Recently, rescoring models have also been reported to increase peptide identifications by integrating the output of multiple SLS methods with other sources of information, such as peptide retention times and sequence features.^26,28^ Usually, the evaluation of SLS approaches is mostly based on the results of the whole SLS pipeline, with no reported independent assessment of the individual pipeline components that are critical for the overall results.

Real-world MS/MS spectra from both data-independent acquisition (DIA) and data-dependent acquisition (DDA) methods often include background noise or uninterpretable peaks,^29^ and can come from a mixture of multiple peptides.^19^ Due to the ubiquitous presence of noise peaks, the design of SLS methods needs to consider how to resist various types and levels of noise while improving identification rates. Recognizing this need, some SLS tools^23^ have sought to test their methods’ robustness against additional noise peaks. However, these peaks are added to query spectra in a randomized way, which tends to be unrealistic since it may place the artificial noise peaks at impossible or highly unlikely masses, thus not accurately representing noise in real-world spectra. It is thus important to evaluate the robustness of peptide SSM scoring functions to more realistic types of noise that may be present in query spectra at different intensity levels in relation to the signal peaks in each query spectrum.

For benchmarking spectral library search, the most widely accepted metric for quantitative evaluation and comparison of SLS methods is the recall/sensitivity at different false discovery rates (FDRs).^8^ The recall represents the number of (presumably correct) SSMs returned by SLS at a fixed FDR (usually 1%), which represents the fraction of returned results that are expected to be incorrect. While some methods have been proposed to estimate error rates using SSM p-value probability that a particular score results from a random match,^20,22^ a more commonly used strategy is to calculate FDR using the Target/Decoy Approach.^30^ Methods for generating decoy spectra^31–33^ currently employ various forms of randomization based on real spectra to maintain realistic spectral profiles. However, it has been observed that different decoy searching methods estimate FDRs in the low FDR range very differently,^21^ and any biases undermining core assumptions for decoy searching may render comparisons between methods invalid or at least questionable,^34,35^ since these biases are model-dependent. Therefore, it is important for an evaluation of SSM scoring functions to be based on a definition of ground truth that is independent from the SLS decoy models.

These limitations in SLS evaluation support the need for an SLS benchmark dataset and an evaluation pipeline that can systematically study the independent impact of preprocessing methods and SSM scoring functions using independently-established ground truth identifications for query spectra, while also testing their resistance to noise and evaluating their performance on real and predicted spectral libraries. We propose new methods to build benchmarks for evaluating the effectiveness and robustness of SSM scoring functions and SLS methods. A benchmark dataset is constructed, including both real and ideal experimental query spectrum sets with additional controllable, realistic noise, and designed to test SSM/SLS performance against real, ideal, and predicted libraries. A baseline SLS pipeline was then established to independently evaluate several widely used preprocessing methods and three standard SSM functions: Cosine, Projected Cosine and Jensen-Shannon divergence. This analysis is then extended to further consider the SSM scoring functions used in SpectraST and EntropyScore, thus showing how the constructed benchmark dataset can assess the performance of critical components used in state-of-the-art spectral library search tools.

## METHODS

### Datasets

Mass spectrometry data from two high-resolution HCD datasets available at MassIVE (http://massive.ucsd.edu) was reused to construct the query sets for the benchmark dataset. Query spectra were obtained from dataset PXD001468 / MSV000079841,^36^ and the source spectra used to generate realistic noise added to the query spectra were obtained from dataset PXD010154 / MSV000083508.^37^ Benchmark spectral libraries of reference spectra based on real experimental data were based on MassIVE Knowledge Base (MassIVE-KB) spectral library version 2.0.15,^7^ obtained from https://massive.ucsd.edu/ProteoSAFe/static/massive-kb-libraries.jsp. MassIVE-KB contains reference spectra for 5,948,126 precursors (3,639,899 unique demodified precursors) of 2,489,522 unique human peptides (i.e., unique sequences regardless of modifications or precursor charge state).

### Query Spectra Set

The Query spectrum set was constructed by re-searching all spectra from PXD001468 / MSV000079841^36^ using MSGF+^5^ implemented in the MSGF-PLUS-AMBIGUITY workflow at MassIVE https://proteomics.ucsd.edu/ProteoSAFe/index.jsp?params={%22workflow%22:%22MSGF-PLUS-AMBIGUITY%22} to search against the UniProt Human reference proteome (downloaded June 2022) with isoforms. The parent mass tolerance was set to 50 ppm and the search required fixed carbamidomethyl on cysteine (C+57.021), and allowed for up to one common sample-handling modification: oxidation of methionine (M+15.995), deamidation (+0.984) on N or Q, pyroglutamate formation (-17.027) on N-term Q, and up to one N-terminal modification: N-terminal acetylation (+42.011) or N-terminal Carbamylation (+43.006).

The resulting Peptide-Spectrum Match (PSM) identifications were filtered at a False Discovery Rate (FDR) of 1% at the spectrum, precursor (i.e., distinct peptide/charge pairs) and protein levels. After calculating all levels of FDR using only the top 1 best-scoring PSM per spectrum, the ambiguity in spectrum identifications was assessed by also retaining runner-up PSMs per spectrum if their spectrum, precursor and protein scores also passed the FDR thresholds established using only the top 1 PSM per spectrum. We emphasize that runner-up PSMs are not used to claim multiple peptide identifications per spectrum – these are only used to establish that a spectrum identification is ambiguous/uncertain in cases where 2+ distinct PSMs could explain the same spectrum at a level passing all FDR thresholds. All spectrum identifications mentioned in the manuscript refer to top 1 identifications unless otherwise noted.

Identified spectra were used to construct the query set by application of filters and other transformations as needed. The precursor MZ of spectra identified through isotopic precursor masses was corrected to the theoretical precursor MZ to eliminate the need to consider precursor isotopes when using these spectra to search against the spectral libraries. Query spectra were also filtered to remove peaks corresponding to immonium ions, as well as peaks annotated as precursor ions based on the identification precursor *MZ* and charge, using a peak-matching tolerance of 0.05 Da.

‘Clean’ query spectra were generated from the real query spectra by retaining only the peaks annotated as ’a’, ’b’, or ’y’ ions, including isotopes and neutral losses of H_2_O or NH_3_, based on their identification (Figure 1A). Peak annotations considered charges 1 and 2 for all ion types, except ’a’ ions, which considered only charge 1. This method for generating a clean spectrum *S*^′^from a given experimental spectrum S is denoted here as *S*^′^ = *SpecAnno(S)*, and was also used to preprocess noise source spectra and library spectra as described below.

**Figure 1.**
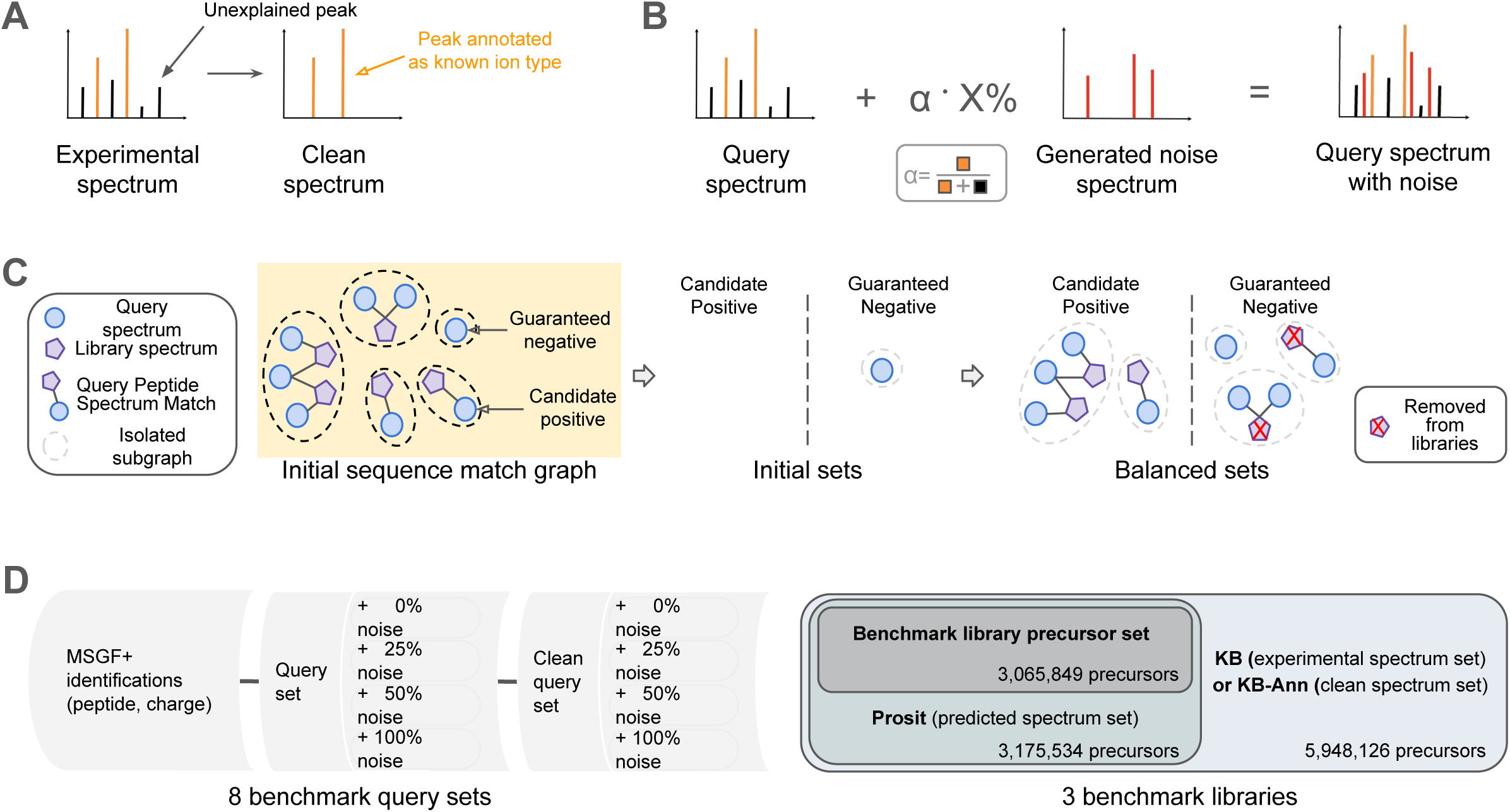
Construction of a Benchmark Dataset. (A) Clean spectra were generated from experimental spectra by retaining only the annotated peaks (orange) of a, b, or y ions including isotopes and H_2_O/NHneutral losses with charges 1 or 2, based on spectrum identifications. (B) Controllable noise is added to a query spectrum *Q* by transferring peaks from a generated noise spectrum *N*. *N* only contains annotated peaks and is generated using another experimental spectrum through the random permutation of the peptide sequence and the corresponding movement of the annotated peaks. *N* is scaled before being added to *Q* so that the total intensity of the added noise peaks equals *X%* of the total intensity of the annotated peaks in *Q*. (C) The sequence-based balancing of candidate library matches is represented as a graph *G*, which contains nodes for query spectra (blue circles) and library spectra (purple pentagons), with edges indicating a sequence and charge state match between the query and library spectra. Query spectra that match to library spectra of the same precursor sequence are considered candidate positives, while those that do not are guaranteed negatives. A sequence match between any two spectra from the same isolated subgraph and a mismatch from different isolated subgraphs are also noted. The candidate positive set is initialized as empty, while the guaranteed negative set is initialized with all guaranteed negative query spectra. Then, each isolated subgraph is considered in descending order of the number of query spectra it contains, and is assigned to the then-smaller set of either candidate positives or guaranteed negatives. Finally, library spectra in the guaranteed negative set are removed from the libraries. (D) The constructed benchmark dataset consists of eight query sets: Each set has 477,803 spectra in both experimental and clean query sets, with additional noise added at noise levels of 0%, 25%, 50%, and 100%. It also includes three spectral libraries: KB, KB-Anno, and Prosit, each containing experimental, clean, and predicted versions of spectra with 3,092,511 shared precursors.

### Adding Noise to Query Spectra

Spectra from dataset PXD010154 / MSV000083508^37^ and their MSGF+ identifications (obtained with the same identification settings described above for query spectra) were used as noise source spectra. As with query spectra, peaks annotated as immonium and precursor ions were filtered out from the noise source spectra. These were then used to generate realistic spectra of randomized sequences whose peaks were added to query spectra to represent different levels of noise. Given a query spectrum *Q*, a noise spectrum *N*′ is generated from a noise source spectrum *N* with a peptide sequence *P*, where *Q* and *N*′ have the same unit precursor *MZ* and cosine similarity < 0.2. In the initial step, *N* is preprocessed to obtain *N* = *SpecAnno*(*WF*_5,50_(*N*)), where *WF*_*K,M*_ is a window filter spectrum preprocessing method where a peak at m/z *m* is retained in the spectrum only if its intensity ranks in the top*-K* most intense peaks in the m/z range [*m – M* Da, *m* + *M* Da]. *N*^′^ is then constructed by generating a shuffled *P’* version of *P* and by moving the peaks annotated in *N* using *P* to the new theoretical m/z values of the same ion fragments in the shuffled sequence *P’*, just as originally described for the generation of decoy spectra in SLS tools.^31^ The resulting *N*′ spectrum was considered suitable for adding noise to *Q* if the cosine similarly (a) between *N* and *N*′ and (b) between *S* and *N*′ were each less than 0.2. If *N*′ was not valid for adding noise to *Q* then a new noise spectrum based on *N* was generated using a new random shuffling of P; if a valid *N’* spectrum was not found after repeating the process up to 10 times then *Q* was removed from the query set. Using α as the percentage of total ion intensity *I* in *Q* that is annotated to known ion types and *X%* as the desired level of noise to be added to *Q*, noise peaks were added to *Q* from a valid noise spectrum *N*′ by scaling their summed peak intensities to α×*X%*×*I* before adding the peaks to *Q* (see Figure 1B), thereby introducing ‘noise’ peaks with valid peptide m/z values and fragmentation patterns at a predetermined fraction of the signal intensity in *Q*. To avoid introducing systematic biases of the same noise source spectrum potentially affecting evaluation results for multiple query spectra, each noise source spectrum was used only once for adding noise to at most one query spectrum.

### Spectral Libraries

Benchmark spectral libraries were designed to be as large as possible while guaranteeing that every spectral library contained reference spectra for the exact same set of precursors. The initial candidate set of precursors was set to the precursors contained in MassIVE-KB (denoted here as the ’KB’ library), after filtering it to retain only high-resolution HCD spectra and to remove all representative spectra originally included from the two datasets used here for query and noise source spectra. Then, a window filter was applied to all reference spectra to keep the peaks whose intensity is ranked in the top-8 most intense peaks in the range of its m/z value ±50 Da. The clean version of MassIVE-KB obtained by applying the *SpecAnno* filter to retain only annotated ion peaks for every reference spectrum is denoted as ‘KB-anno’. The precursor sequences and charge states were used to generate predicted reference spectra using the online Prosit^11^ platform at https://www.proteomicsdb.org/prosit, which only supports predictions for sequences of length 7 to 30 amino acids long and only supporting two types of modifications (carbamidomethyl on cysteine and oxidation of methionine). Prediction parameters were set to recommended values^11^ using the Prosit_2020_intensity_hcd model with a fixed collision energy of 30. Prosit predicted spectra set each precursor *MZ* to its theoretical value, and MS/MS peaks with predicted intensities are reported at the theoretical *MZ* of b/y ions with charge states up to the precursor charge. Peaks with negative predicted intensity values were removed from the predicted spectrum, and a predicted spectrum was removed from the predicted spectra set if there were no peaks after filtering out peaks with negative intensities. Precursors that could not be successfully predicted using Prosit were also removed from the KB and KB-anno benchmark libraries.

### Precursor Selection

Precursors selected for inclusion in the benchmark query and spectral library sets were categorized as either (a) *candidate positives*, corresponding to query precursors for which the libraries contain a reference spectrum for the exact same precursor (with allowance for amino acid "I" matching "L" and different site localizations for oxidation of methionine), or (b) *guaranteed negatives*, corresponding to query precursors where the library is guaranteed to not contain a reference spectrum of the same precursor. This categorization is solely based on matching precursor sequences and charge states, with no consideration for the level of similarity between the spectra identified to the precursors. Enforcement of this categorization required the exclusion of precursors that could not be unambiguously assigned to either category. Two peptide sequences are defined to be an *ambiguous match* if (1) they have the same length, (2) their sequences are not exactly the same and (3) they differ by at most one amino acid pair ("Q", "K"), ("Q", "E") or ("N", "D") after replacing all instances of "I" with "L" and ignoring deamidation on N or Q in both strings.

For a query spectrum to qualify as a query set precursor, it must meet the following criteria: (1) it should be identified by MSGF+ as described above and have no tie for its best-scoring identification, (2) it should have been paired with a noise spectrum for noise addition, (3) its precursor identification should not have any deamidation, since these are often confused with ^13^C carbon isotopes, (4) if its top 1 identification is not an exact match to the library, then (a) the top 1 identification should not be an ambiguous match to the library and (b) none of its runner-up identifications (if any) should be an exact or ambiguous match to the library, (5) there should be at least one library spectrum that has the same precursor charge state and a precursor *m/z* value within the range of is precursor *m/z* value ±0.05 Da.

Given the high overlap between the initial spectral libraries (with a comprehensive coverage of millions of human precursors) and the input query set (with mostly commonly-detected human precursors), the initial number of candidate-positive query spectra/precursors was much higher than the number of guaranteed-negative spectra/precursors. As such, it was necessary to rebalance these sets by removing a subset of precursors from the library, thus converting candidate-positive query spectra into guaranteed-negative query spectra (since the correct match is removed from the benchmark library). The selection of which precursors to remove from the library was made using a *sequence match graph* (Figure 1C) defined as follows: a sequence match graph *G* = {*V*, *E*}, is defined by a set of nodes *V* = {*V*_*q*_, *V*_*l*_} that represent all query (*V*_*q*_) and library (*V*_*l*_) spectra, with edges added between a query spectrum node and a library spectrum node with the same charge state and an exact sequence match between any of the identifications from each node. As such, query spectrum nodes connected to any library spectrum node represent *candidate positives*, while those that are not connected to spectrum library nodes represent *guaranteed negatives*. *Precursor clusters* are the isolated subgraphs in *G*, comprising connected nodes, with no edges between nodes from two distinct isolated subgraphs. Isolated library spectrum nodes are removed from *G*.

To balance the number of spectra and precursor clusters of candidate positives and guaranteed negatives, a greedy method was introduced to assign precursor clusters to new candidate positive and guaranteed negative sets (Supplementary Algorithm S2). Specifically, the candidate positive set is initialized as empty, while the guaranteed negative set is initialized with all query spectra that do not have a top 1 identification that is an exact match to the library. Then, all isolated subgraphs are traversed in descending order of their number of query spectra, and assigned to the then-smaller set of either candidate-positives or guaranteed-negatives. Library spectra in the guaranteed negative set are then removed from the selected library precursor set (Figure 1C).

The real and clean query spectrum sets share the same set of selected query precursors. Additionally, for each experimental spectrum in the query set and its clean version in the clean query set, the peaks from the same noise spectrum are used to add noise at levels of 25%, 50%, and 100% of the annotated peaks intensity in the query spectrum. Based on the selected library precursor set, three libraries are used for the benchmark: KB with MassIVE-KB experimental reference spectra, KB-Anno with clean MassIVE-KB experimental reference spectra, and Prosit with predicted spectra. (Figure 1D)

### Benchmarking Spectrum-Spectrum Matching Scoring Methods

Spectrum preprocessing and spectrum-spectrum matching (SSM) are two critical steps in spectral library searches (SLS). Therefore, we built a baseline pipeline using widely accepted methods from these two steps to form standard scoring functions. Five standard preprocessing methods were considered: (1) square root intensity transformation, (2) window filter, (3) signal-to-noise ratio filter, (4) top peaks filter and (5) high-intensity filter; as well as three standard SSM scoring functions: Cosine (Cos), Projected-Cosine (ProjCos), and Jensen-Shannon divergence (JS-divergence). Detailed definitions of these considered methods are provided in Supplementary Text S1.

The scoring functions used in current SLS methods can incorporate more complex preprocessing steps and can include postprocessing steps to renormalize spectrum matching scores with additional information beyond just two spectra. As such, the SLS scoring functions used in two state-of-the-art SLS methods, SpectraST^21^ and EntropyScore,^23^ are also benchmarked here. SpectraST^21^ is one of the most popular spectral library search engines, and EntropyScore^23^ was initially proposed for identifying small molecules, but it has recently shown promise in peptide identification. Supplementary Text S2 provides additional information on the differences between these two methods and standard SLS scoring functions.

The implementation of JS-divergence and EntropyScore is modified based on the official code repository (https://github.com/YuanyueLi/SpectralEntropy) to allow spectrum combination based on matched peaks within m/z tolerance and additionally output the second highest-scoring matched library spectrum whose sequence does not match the highest-scoring match to the same query spectrum. SpectraST version v5.0 was used as provided in the Trans-Proteomic Pipeline^38^ (http://www.tppms.org/).

### Benchmark Settings and Metrics

All library searches use a precursor m/z tolerance of 0.05 Da and a peak m/z tolerance of 0.05 Da Query spectra can only match library spectra with the same precursor charge state. Spectrum matches are evaluated by comparing the identification of the top-scoring library spectra with the top 1 MSGF+ identification for a given query spectrum. A query spectrum match is *correct* for a given threshold *T* if its top-scoring library spectrum match has a score ≥ *T*. In cases of tied top scores, at least one match has an exact sequence match to the query. The precision and recall are computed as:

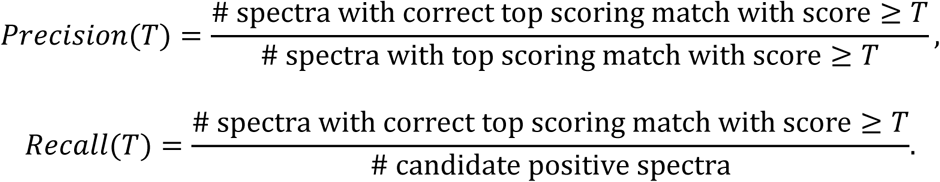

Due to the ambiguity that exists in spectrum identification caused by (a) possibly multiple MSGF+ identifications for the same query spectrum and (b) possibly more than one library spectrum passing a given threshold *T*, we define a query spectrum match as *ambiguous correct* for threshold *T* if either of its top-2 scoring library spectrum matches (1) has a match score ≥ *T* and (2) has an exact sequence match to the query identification or any of its runner-up identifications. Then, the ambiguous precision and recall are computed as

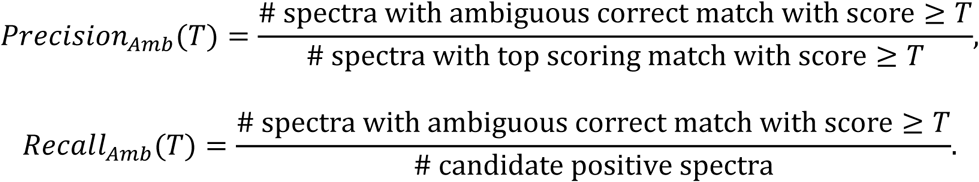

## RESULTS AND DISCUSSION

### Benchmark Dataset Statistics

In a typical peptide spectral library search, experimental spectra (i.e., query spectra) are compared to a *peptide spectral library*, a collection of reference spectra that were previously confidently identified or that were obtained from known peptides (e.g., by peptide synthesis). By comparing the query spectra with the reference spectra in the library using similarity scores computed by spectral library search (SLS) methods, the peptide sequence that best matches the query spectra can be identified. A benchmark dataset that contains eight query sets was constructed to assess the effectiveness and robustness of the standard and state-of-the-art SLS methods on various libraries: 476,063 spectra in each experimental and clean query sets, with additional noise added at the noise levels of 0%, 25%, 50%, and 100%. The benchmark also considered three spectral libraries: KB, KB-Anno, and Prosit, containing experimental, clean, and predicted versions (correspondingly) of spectra for 3,065,819 precursors (see Figure 1D). The precursor m/z distribution in query sets and spectral libraries (Supplementary Figure S1), shows that the search space for most query spectra is large, with a maximum of ∼850 possible library spectra as matching candidates within half of the precursor m/z tolerance (0.05 Da), while the size of the search space size can vary significantly across query spectra. Spectra for precursors with charge 2 are the most common, slightly higher than the number of spectra for charge 3 precursors in the library but about double in the query set. Precursors of varying charge states exhibited very similar precursor m/z distributions in both candidate positive and guaranteed negative sets (see Figure S-1C).

As shown in Figure 2A, most query spectra have substantially more peaks than library spectra, but a similar averaged number of b/y ions peaks with a charge state of 1 or 2 as KB and KB-Anno. In contrast, the Prosit model tends to over-predict b/y ion peaks - the predicted library has about twice as many b/y ion peaks as both query spectra and KB/KB-Anno library spectra on average. The clean query spectra have ∼50 peaks on average while query spectra have ∼193 peaks on average, thus reflecting the typical observation that most spectrum peaks cannot be assigned to common fragment ions of the peptide sequences assigned to each spectrum. Added noise peaks add an average of ∼26 peaks to each spectrum, which only rarely (and randomly) match b/y ions of the assigned query identification. Figure 2B shows that the averaged explained intensity (EI) of KB spectra (70%) is lower than that of KB-Anno and Prosit (both 100% EI), but better than that of query spectra (55%). Note that spectra in the query set and KB contain unannotated peaks, whereas KB-Anno and Prosit do not. The clean query spectra (containing only annotated peaks) also have 100% EI so searching the clean query set simulates the best-case scenario of "ideal" experimental query spectra with all the typically-generated signal but with no noise. Adding X% noise to both clean query and query spectra simulates the case of spectra with an additional co-eluting peptide which does not match the spectral library and whose peaks have X% as much intensity as the summed intensity of the peaks annotated by the MSGF+ identified peptide (see Fig 1B).

**Figure 2.**
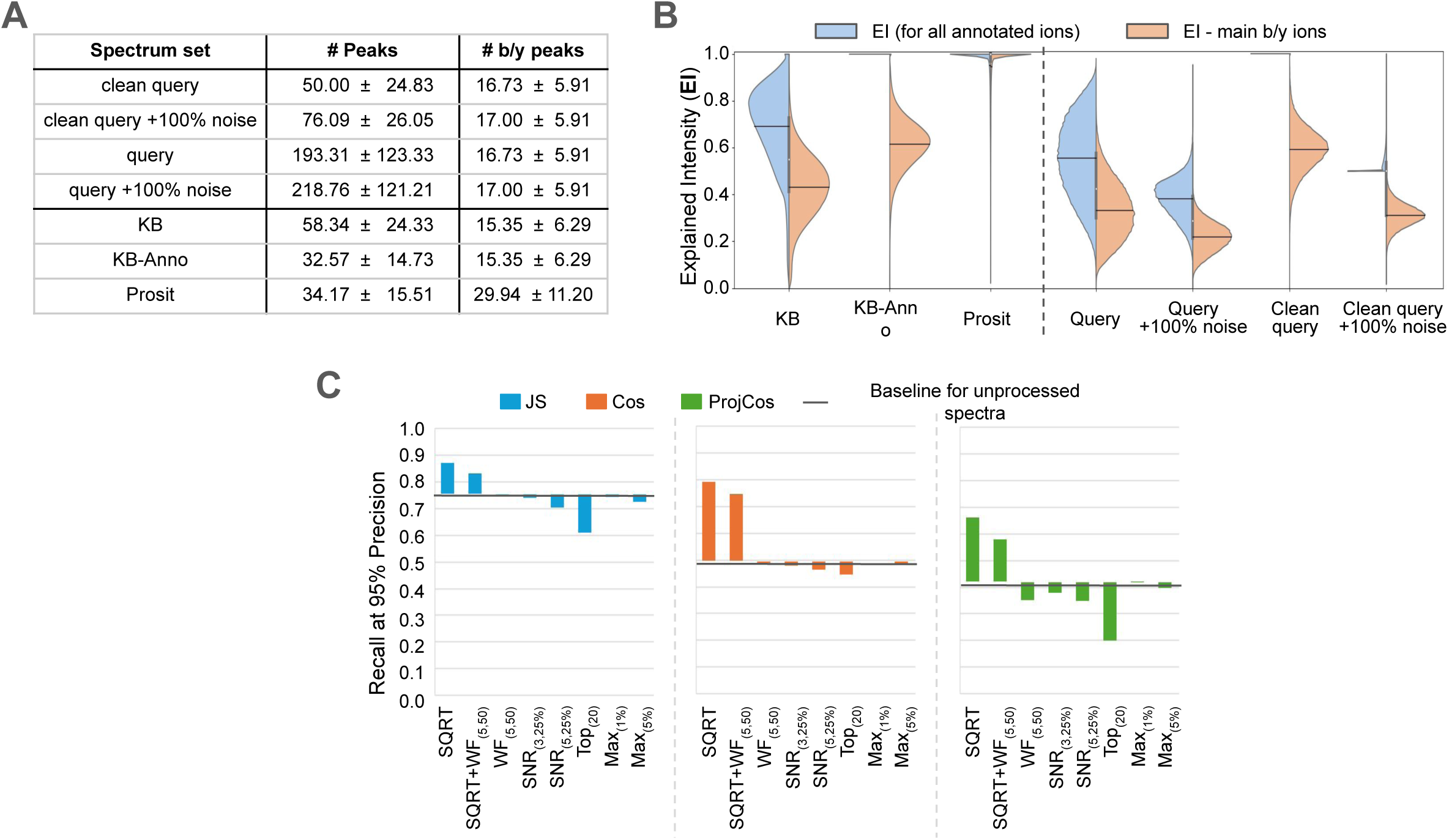
Benchmark dataset statistics and the impact of preprocessing methods. (A) The mean and standard deviation of peaks, along with the peaks annotated as main b/y ions with either a charge of 1 or 2. (B) The distribution of explained intensity (EI) for annotated ions, as shown in Figure 1A, and the main b/y ions with either a charge of 1 or 2. (C) The changes in recall at 95% precision after applying different preprocessing methods or their combinations for JS-divergence (blue), Cosine (orange) and Projected-cosine (green).

### Impact of Preprocessing Methods

To evaluate the performance of various types of spectrum preprocessing, we generated 27 different scoring methods by combining the 3 standard SSM functions (Cos, ProjCos and JS-divergence) with 9 preprocessing scenarios for query spectra: (1) no preprocessing, (2) *SQRT*, (3) *SQRT* + *WF*_5,50_, (4) *WF*_5,50_, (5) *SNR*_3,25%_, (6) *SNR*_5,25%_, (7) *TOP*_20_, (8) *MAX*_1%_, and (9) *MAX*_5%_.

We first evaluate the impact of these preprocessing scenarios in the typical search scenario - searching experimental query spectra against an experimental spectral library. Figure 2C demonstrates the changes in recall at 95% precision when switching from unprocessed spectra to spectra processed in eight preprocessing scenarios. The square root transformation, SQRT, consistently improves all SSM functions the most. It indicates that all SSM scoring functions put too much emphasis on high-intensity peaks and confirms that reducing the relative importance of intense peaks can improve the inner-product- and entropy-based SLS methods.^8,23^ The results of Cos and JS-divergence also show that filtering out peaks often leads to worse performance, even though it increases the signal-to-noise ratio. A possible reason for this could be that signal peaks are being removed along with noise peaks and causing a worse aggregate effect than the benefits accrued from the noise reduction and increase in signal-to-noise ratio. The impacts of these spectrum preprocessing methods are similar for Cos and JS-divergence at 99% precision, although the baseline is significantly lower (see Figure S-2A). The exception was ProjCos whose recall at 99% was too low to consider when comparing preprocessing methods (the performance of ProjCos is further discussed below).

Considering the impact of these preprocessing methods when searching against the predicted Prosit library (see Figure S-2C and S-2D), shows that the combined use of SQRT and *WF*_5,50_ yields the highest increase in recall at both 99% and 95% precision of all SSM functions, with just SQRT usually yielding almost the same gains in recall.

Overall, SQRT and the combined use of SQRT and *WF*_5,50_ were the two most effective preprocessing methods when searching experimental query spectra against both experimental and predicted libraries for all standard SSM functions. For all results presented below, SQRT preprocessing was applied to all spectra before using any standard SSM functions.

### Exploring upper bound on search performance

Clean query spectra containing only peaks annotated as a/b/y-ions (including isotopes and neutral losses) represent the best-possible quality (i.e., ideal), experimental query spectra. These clean query spectra were searched against all benchmark libraries to evaluate and compare the standard SSM scoring functions (Cos, ProjCos and JS-divergence) with the two scoring functions used in the state-of-the-art SLS methods (SpectraST^21^ and EntropyScore^23^) in the best-case scenario where search FDR would be properly controlled for a 1% error rate.

The results (shown as black horizontal lines in Figure 3A) fall into three distinct tiers. The top tier includes SpectraST and JS-divergence, with SpectraST presenting the overall best-performing SSM score. ProjCos defines the bottom tier for reasons that will be discussed in more detail later in this section. The middle tier consists of EntropyScore and Cosine, with Cosine (the most widely used scoring function) performing the worst across almost all scenarios. Within the middle tier, Cosine’s performance was surprisingly poor (with a recall of just 57%) when searching against KB. Conversely, EntropyScore performed slightly better than Cosine, except for predicted spectra (Prosit), which aligns with previous reports on metabolomics spectra.^23^ Both scoring functions yielded better results against the KB-anno and Prosit libraries, which is expected since the clean query spectra contain peaks only for the same ion types that were used to construct KB-anno from KB (mainly b/y-ions) which are also mostly expected to match Prosit-predicted ions. The best results for the top tier were also achieved with KB-anno, with 95% recall for SpectraST and 89% recall for JS, which is also JS’s most competitive scenario vs SpectraST.

**Figure 3.**
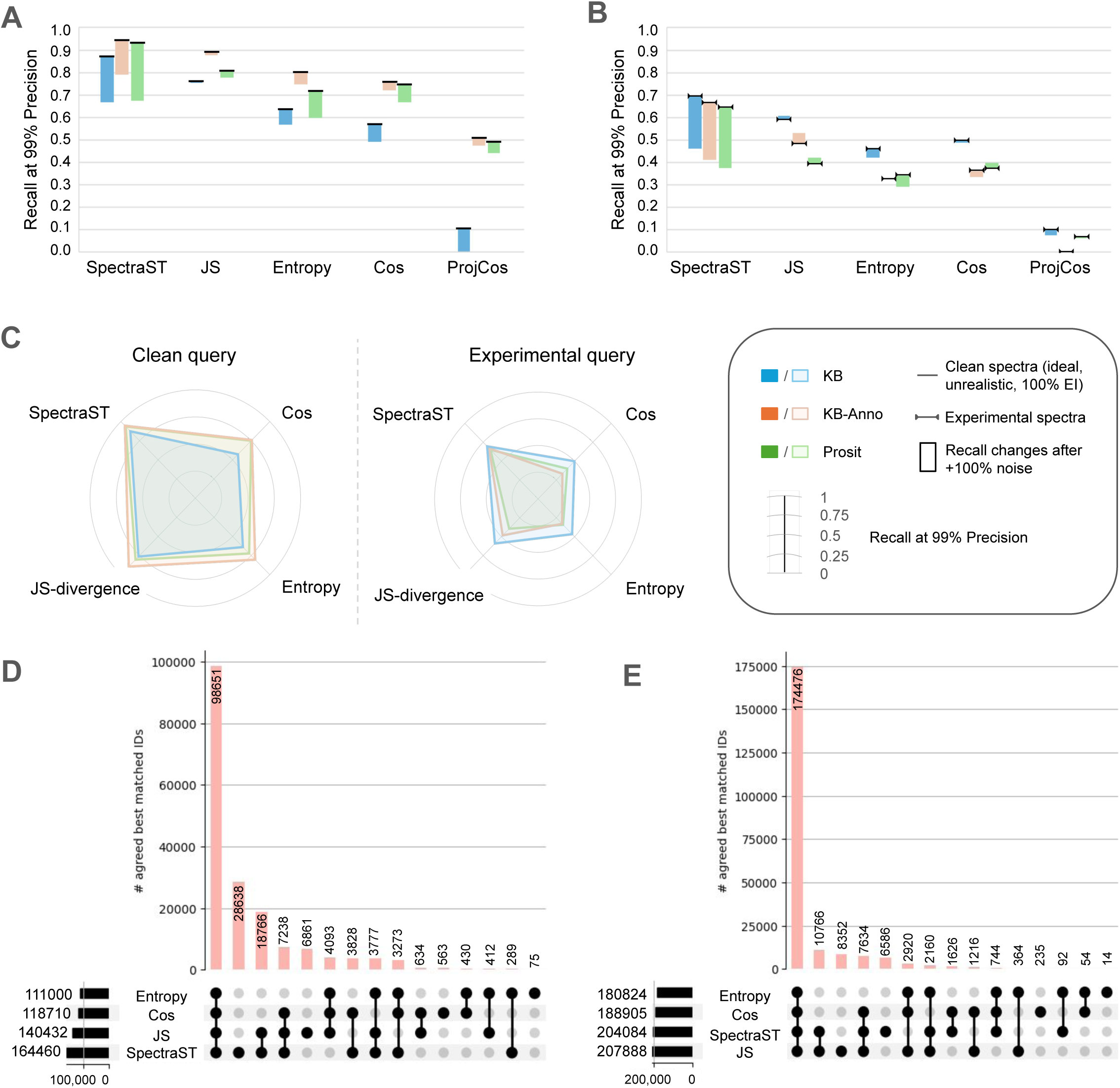
Results on searching “ideal” and real experimental spectra with various levels of additional noise. (A) Recall at 99% precision when searching clean query set, i.e. “ideal” experimental spectra without and with additional 100% noise. (B) Recall at 99% precision when searching query set, i.e. “real” experimental spectra, with additional 0% and 100% noise. (C) Library comparison on recall at 99% precision when searching clean query (left) and experimental query (right). (D) Venn diagram of the number of correct top-scoring spectrum matches from Entropy, Cos, JS and SpectraST when searching query set against KB at 99% precision respectively. (E) the same Venn diagram but at 95% precision.

Overall, the clean/ideal spectra search results indicate that the spectrum library search problem is not completely solved, even in the unrealistic best-case scenario of completely noise-free spectra, even though the best-performing scoring function (SpectraST) can perform quite well if paired with either the KB-anno (95% recall) or Prosit (94% recall) spectral libraries.

### Search "ideal" spectra with known noise fractions

The next round of experiments was designed to evaluate the performance of the SSM scoring functions when the spectra contain a known fraction of noise, which was defined as peak ion intensity that was not derived from any fragmentation of the peptide that generated the query spectrum (see Methods for details of how these were generated). In brief, when adding *X*% noise to the clean query spectra, the fraction of noise is 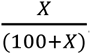. While noise was added at different levels to study the progression of its impact on the search results (see Figure S3 for the full set of results), the results shown in Figure 3A and discussed here consider only the case of adding 100% noise (which means the spectrum contains as much signal peak intensity as it contains noise peak intensity), since this scenario is representative of what would generally be considered a high-quality DDA spectrum identification with approximately 50% explained peak intensity (EI).

The most striking result from this experiment was the very significant reversal of top performers, with JS now performing substantially better than SpectraST across all libraries. While the performance of SpectraST dropped more than 10-20 percentage points (depending on the library), JS was found to be nearly unaffected by the noise added to query spectra and retained a top performance of 87% recall when paired with KB-anno. Another important observation was that searches against KB-anno were consistently more tolerant to the noise added to the clean query spectra than searches against predicted spectra. Given the high number of ions predicted by Prosit (see Figure 2A), one possible explanation for this observed difference between the libraries could be due to KB-anno containing only signal peaks that were observed before (and are thus most likely to be observed again). In contrast, predicted spectra containing over-predicted ion peaks may have caused a higher rate of false matches to noise peaks, which could increase the number of false positive SSMs and thus reduce overall sensitivity at the same fixed error rate.

### Results on typical query spectra

When searching real query spectra against all libraries, all scoring functions performed quite substantially worse than they had on clean query spectra, and even significantly worse when searching real query spectra with added noise (see Figure 3B). Inspection of the search results suggested that a possible explanation for this observation is that real query spectra contain unannotated peaks that are more likely to cause false positive matches to the library. In particular, cases such as those illustrated in Supplementary Figure S-4 show that some query spectra actually contain peaks from multiple peptides, which could potentially cause different methods to select different top matches for the same query spectrum depending on whether the scoring function emphasizes the fraction of matched intensity or the number of matching peaks.

SpectraST remained the best-performing search tool and JS-divergence remained the second-best scoring function, with the gap in recall rates between the two remaining at ∼10 percentage points. However, just as had been observed when adding noise to clean query spectra, the performance of SpectraST dropped by 20+ percentage points when noise was added to the real query spectra, which now simulates the common situation where experimental spectra contain peaks from 2+ co-eluting peptides. Also similar to the results on clean query spectra, Figure 3B shows that JS-divergence was essentially unaffected by the addition of noise peaks and performed significantly better than SpectraST in this important scenario due to its higher tolerance for noise peaks in query spectra.

In contrast to the results on clean query spectra, EntropyScore performed slightly worse than Cosine on real query spectra. However, as the results were similar across most libraries, this suggests that neither scoring function is likely to significantly outperform the other in general. A clearer finding from these results is that JS-divergence consistently outperformed EntropyScore, usually by achieving recall rates higher by more than 10 percentage points. This indicates that the EntropyScore’s normalization of peak intensities does not benefit peptide spectra as it does for metabolomics spectra. JS-divergence was also consistently better than Cosine, and oftentimes quite significantly so - especially when it achieved its best overall performance of searching real spectra against the original MassIVE-KB library.

Overall, the observed search performance was generally poor, with observed recall rates of only 70% for the best-case scenario (SpectraST paired with KB or KB-anno spectral libraries) and with an even worse recall rate of 61% for only JS paired with KB when searching query spectra with added noise simulating the presence of noise peaks or peaks from co-eluting peptides.

### Comparison of spectral library performance

Library quality is a critical factor influencing the performance of spectral library search. We compared the performance of SLS methods using "ideal" and "typical" experimental spectra against experimental, clean, and predicted benchmark libraries. Results illustrated in Figure 2C clearly showed that real libraries with experimental spectra (KB and KB-Anno) outperformed the predicted library (Prosit). However, the predicted library also performs very well when real libraries are unavailable.

When clean query spectra were searched, KB performed the worst in all cases but this was not surprising given that these query spectra were unrealistically devoid of unannotated peaks, even though a significant fraction of these peaks in KB is likely to correspond to unannotated peptide fragmentation events (e.g., from internal fragments resulting from multiple peptide fragmentation events). However, the KB-anno filtered version of KB performed the best across all libraries for all scoring functions, thus indicating that libraries derived from experimental spectra still provide reference spectra that are more similar to query spectra than those obtained with the Prosit prediction model. These results also highlighted two points regarding predicted spectra. First, SpectraST library search of the clean/ideal query spectra using predicted spectra still achieved nearly optimal performance, again indicating that predicted spectra can be a valuable resource when experimental spectral libraries are not readily available. However, the second noteworthy observation is that this competitiveness only holds if predicted spectral libraries are paired with a "compatible" scoring function such as SpectraST or Cosine (both cosine-based scoring functions), while the comparative performance is significantly worse when used for JS-divergence or EntropyScore (both probability-based scoring functions).

Finally, the experimental query set search results revealed MassIVE-KB as the best library for searching real experimental query spectra, often by a significant margin; for example, the recall was higher by over 20 percentage points when searching KB vs the Prosit-predicted library when using JS-divergence as the SSM scoring function. Since the results using KB were almost always also substantially better than when using KB-anno, a possible explanation for this higher performance could be that the unannotated peaks in the query spectra match the unannotated peaks in library spectra, which were present in MassIVE-KB reference spectra because these were selected from large collections of experimental data, but filtered out in KB-anno because it retained only annotated ion peaks.

### Identification agreement across methods

Beyond comparing the recall rates for the various SLS methods, the more detailed analysis of identifications reported in Figure 3D shows that different methods emphasize the identification of different peptides. Although SpectraST was the most sensitive approach, identifying a total of 164,460 spectra at 99% precision, there were 13,068 other spectra (7.3% of all 177,528 identified spectra at this level of precision by at least one approach) that were correctly identified by other SLS methods but were missed by SpectraST.

Furthermore, the comparison of identification results at 95% precision, as shown in Figure 3E, shows a significant increase in the overlap of identifications obtained by all approaches, especially the 95% overlap between JS-divergence and SpectraST identifications (which was substantially higher than the 74% overlap observed at 99% precision). This suggests that most of the differences between the methods result from minor variations in the ranking of the top-scoring SSMs, causing different true positive matches to fall below the score threshold for 99% precision but still score high enough to overlap at 95% precision.

A noteworthy finding from these results is that JS-divergence becomes the best at 95% precision, where its recall rate of 87% is significantly higher than its own (59%) or SpectraST’s (70%) recall rates at 99% precision. This suggests that JS could provide substantially better results if it was able to better distinguish between true matches and a small number of false positive matches in this high-scoring range of spectrum-spectrum matches.

### Results on Projected-Cosine

While ProjCos is a scoring function primarily used for analyzing spectra of co-eluting peptides, it also needs to properly process the simplest possible case of spectra that contain fragment ions from just one single peptide. However, as shown in Figures 3A and 3B, ProjCos performed dramatically worse than all other scoring functions across all libraries. A closer inspection of the top-scoring ProjCos matches revealed two important factors related to its poor performance. Firstly, not all of its high-scoring matches marked as FPs should be marked as such - there are instances where the query spectrum really is a chimeric spectrum containing fragments from two different peptides, and ProjCos favors a different top-scoring identification than the one selected by the database search engine MSGF+ for the same spectrum. However, this was observed for only a small fraction of the high-scoring matches marked as FP. The second, and much more important factor, was that artificially high-scoring matches were caused by matching very few high-intensity peaks and ignoring most fragment ions (see Figure 4A). Therefore, this substantially increases the likelihood of false positive matches since it does not penalize unmatched query peaks like Cos and other functions do.

**Figure 4.**
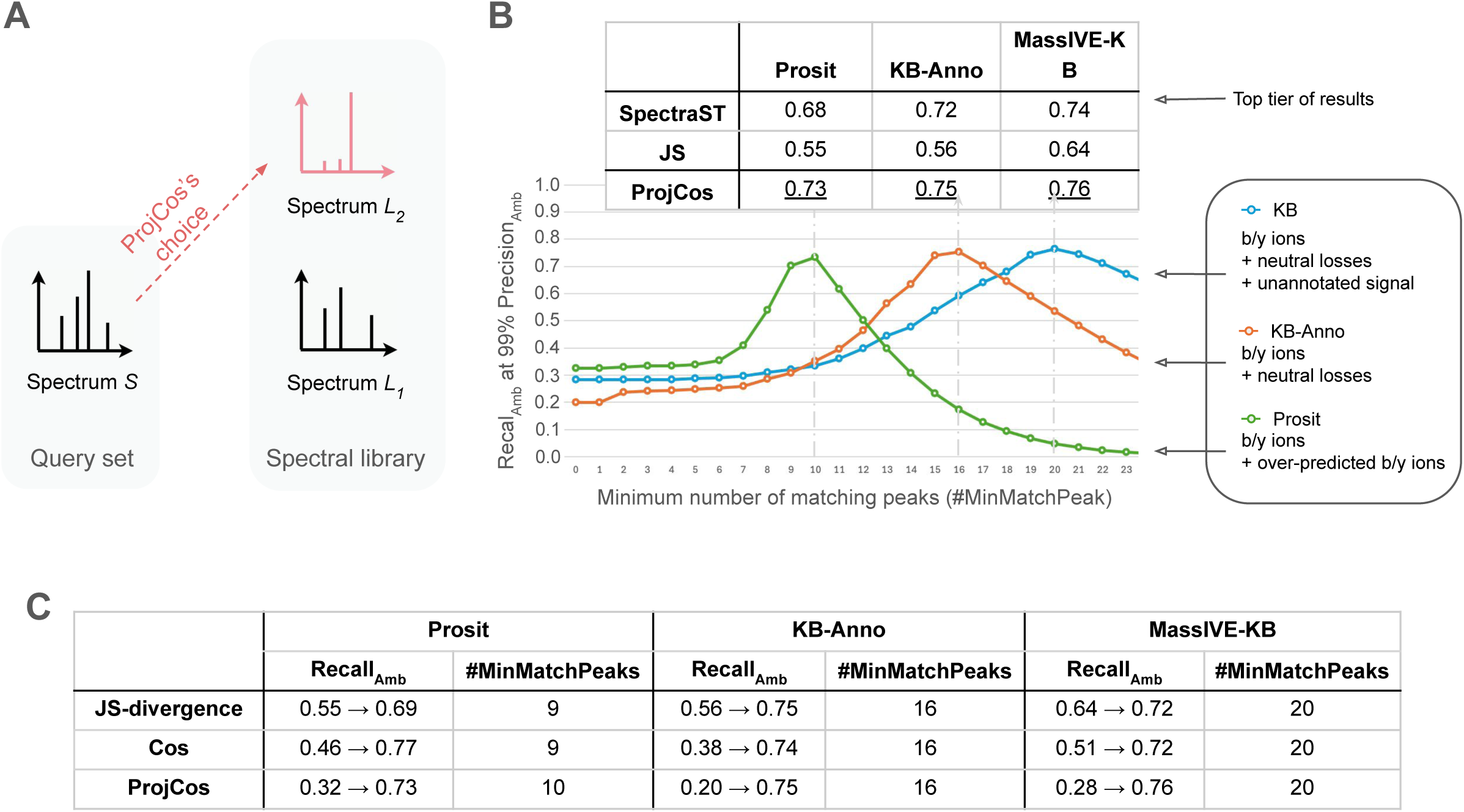
Results on ProjCos and the impact of the minimum number of matching peaks (#MinMatchPeaks). (A) ProjCos tends to assign library spectra with fewer peaks higher score for a given query spectrum and ignore most fragment ions. (B) The ambiguous recalls at 99% ambiguous precision vary dramatically depending on the minimum number of matching peaks. The performance can extremely poor recall (as low as 0%) with no matching peak requirements, to consistently outperforming all other methods when the best-possible value for the required minimum number of matching peaks is selected for all libraries. (C) The changes of ambiguous recall at 99% ambiguous precision from without requiring the minimum number of matching peaks to the ones that achieve the optimal performance for scoring methods JS, Cos and ProjCos with additional use *WF_5,50_* on query spectra.

Following these observations, we further considered how the performance of ProjCos would be affected by requiring a minimum number of matched peaks between two spectra. As shown in Figure 4B, the performance of ProjCos with additionally applying the window filter *WF*_5,50_ on query spectra can vary dramatically depending on the minimum number of matching peaks required when comparing a query spectrum to a reference spectrum. The performance varies from extremely poor ambiguous recall (the recall based on top 2 library matches and the top K MSGF+ identifications, see details in Methods) as low as 20% with no matching peak requirements, to consistently outperforming all other methods when the best possible value for the required minimum number of matching peaks is selected for all libraries. Given the disproportionate importance of this parameter value, we further studied its impact on the Cosine and JS-divergence scoring functions as well and found (see Figure 4C) that the performance of all scoring functions becomes very similar when the optimal minimum number of matching peaks is properly selected.

Two important observations were also noted across all experiments. First, the minimum number of matching peaks that worked best for each library was consistently smallest for Prosit and largest for KB. This was as expected since the required number of matched peaks reduces to a required number of matched b/y ions for predicted spectra (since they only contain monoisotopic b/y ions), which is a much stronger requirement than just a number of matching peaks regardless of ion type. Optimal matches to KB required the highest minimum number of matching peaks because its spectra contain the largest number of peaks - they contain all annotated ions (i.e., main b/y ions, their isotopes and neutral losses) as well as unannotated peaks of significant intensity in the reference spectrum selected for each precursor.^5^ The number of peaks required for matches to KB-anno was found to be in between the other two libraries, which was also as expected since this library contains peaks annotated b/y ions as well as their isotopes and neutral losses, so it needs a higher minimum number of matching peaks to enforce a number of matched b/y-ions similar to what ends up being applied for Prosit-predicted spectra. Conversely, it does not contain the unannotated peaks present in the KB library, so it does not require as many matching peaks as when matching to KB to achieve the same level of significant results. The second important observation was that the optimal performance of each scoring function was highly dependent on the specific selection of the minimum number of matching peaks. While all scoring functions achieved comparable near-optimal performance with the precise setting of this parameter value, their performance also degraded relatively quickly with small variations in the choice of values for this parameter. As such, these results indicate that the performance of the scoring functions may depend quite substantially on the peak statistics of different mass spectrometry runs, which should be taken into consideration by any methods based on the scoring functions evaluated here.

## CONCLUSION

This paper proposes new methods to build benchmarks for Spectrum-Spectrum Matching (SSM) scoring functions, which are the most important component of peptide identification algorithms for spectral library search. These methods were then used to build a benchmark dataset to evaluate the performance of the most commonly used SSM scoring functions: Cos, ProjCos and JS-divergence, as well as the impact of 9 common spectrum preprocessing scenarios on each of them. Then, 2 additional scoring functions used in state-of-the-art spectral library search tools (SpectraST^21^ and EntropyScore^23^) were also evaluated.

The results revealed that the Cosine scoring function, despite being the most used SSM scoring function, performed significantly worse than probabilistic (JS-divergence) or mean-corrected (SpectraST’s per-spectrum cosine F-score) scoring functions, with SpectraST performing best for spectra with little-to-no noise, while JS-divergence was the most resistant to noise across all libraries. The requirement of a minimum number of matching peaks was identified to have a significant influence on the performance of all standard SSM functions (Cos, ProjCos and JS-divergence), with ideal settings enabling the standard SLS methods to be competitive with or even outperform SpectraST. Notably, even minor variations in the choice of the required minimum number of matching peaks could cause significant fluctuations in recall.

Moreover, these results also raise serious concerns with approaches allowing identifications from very few peaks. It was observed that ProjCos, which is widely used for searching multi-peptide spectra, requires at least 10+ matching peaks to achieve reasonable levels of recall at 99% precision, calling into question the error rates of methods that use similar scoring functions for detection of peptides in DIA mass spectrometry runs while requiring no more than 2 matching peaks.^27^

By having established this preliminary benchmark for the comparison of SSM scoring function, it is anticipated that the same or similar methods should be suitable for establishing follow-up benchmark datasets for assessing library searching of spectra from different types of peptide fragmentation modes (such as CID, ETD, etc.) or for evaluating rescoring methods used in SLS methods (such as PeptideProphet^39^ and Percolator^15^). Additionally, these methods can be used to construct benchmark datasets with alternative ground truth definitions, such as using search results from a search engine other than MSGF+^5^.

## ASSOCIATED CONTENT

### Data Availability Statement

All benchmark datasets, libraries and spectrum level identification results are available in the MassIVE repository with identifiers MSV000095946 and PXD056205.

### Supporting Information

Supplementary Text S1: Baseline pipeline for standard SLS scoring methods. Supplementary Text S2: Differences between SpectraST/EntropyScore and standard SLS scoring functions.

Supplementary Algorithm S1. A greedy method for finding the pairing PM between two spectra. Supplementary Algorithm S2. Balancing candidate positives and guaranteed negatives.

Supplementary Figure S1. Precursor Mass/Charge distribution per charge state. Supplementary Figure S2. Additional results on the impact of preprocessing methods.

Supplementary Figure S3. Additional results on the Impact of noise.

Supplementary Figure S4. An example of the query spectrum from multiple peptides\, which causes different methods to select different top matches.

## ACKNOWLEDGMENTS

We thank Pisit Wajanasara and Jeremy Carver for helpful discussions during the project development. This work was partially supported by the National Institutes of Health (grants NLM R01LM013115, NIGMS R24GM148372) and National Science Foundation (grant ABI1759980).

## Supporting Information

### Supplementary Text S1: Baseline pipeline for standard spectrum-spectrum matching scoring methods

A baseline pipeline is proposed, using standard methods in spectrum preprocessing and spectrum-spectrum matching (SSM), to evaluate standard spectral library search (SLS) scoring functions. The evaluated spectrum preprocessing methods and Spectrum-Spectrum Match (SSM) scoring functions required implementing methods using the following definitions.

#### Supplementary Definition 1

A *spectrum* is defined as a set of *peaks*, i.e., (mass-over-charge or m/z, intensity). Specifically, we represent a spectrum *S* = {(*MZ*_*i*_, *I*_*i*_)}_*N*_, where *i* = 1, 2, …, *N*, where *N* is the number of peaks in *S*. Here, *s*_*i*_ = (*MZ*_*i*_, *I*_*i*_) is the *i* −th peak of the spectrum, and *MZ*_*i*_ and *I*_*i*_ refer to the mass-over-charge ratio (m/z) and the intensity of the i-th peak.

#### Supplementary Definition 2

Given a peak *m/z* tolerance *T*, two peaks *s*_*i*_ = (*MZ*_*i*_, *I*_*i*_) and *s*_*j*_ = (*MZ*_*j*_, *I*_*j*_) *match* if |*MZ*_*i*_ − *MZ*_*j*_| ≤ *T*. Thus, a peak matching function is defined as

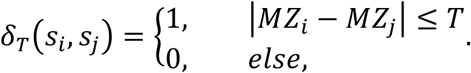

#### Supplementary Definition 3

A *spectrum-spectrum match (SSM) scoring function* takes two tandem mass spectra as input and outputs a score (usually ranging from 0 to 1) to describe the similarity between these two spectra. A higher score indicates a more significant similarity between the two given spectra.

Formally, a preprocessing method can be regarded as a function that takes a mass spectrum as input and outputs another spectrum. Five standard preprocessing methods were considered:

1) Square root transformation, 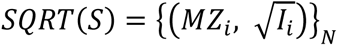 for all *s*_*i*_ ∈ *S*.
2) Window filter, *WF*_*K,M*_(*S*), keeps the *j*th peak if its intensity *I*_*j*_ has rank ≤ *K* (where rank 1 is the highest intensity peak) for all spectrum peaks with MZ in [*MZ*_*j*_ − *M*, *MZ*_*j*_ + *M*].
3) Signal-to-noise ratio filter, *SNR*_*K,X*%_(*S*), keeps the peaks whose intensity is *K* times higher than the average intensity of the *X*% lowest-intensity peaks (across the whole spectrum *S*).
4) Top peak filter, *TOP*_*K*_(*S*), keeps the *K* highest intensity peaks of *S*.
5) High-intensity filter, *MAX*_*K*%_(*S*), where *K*% ∈ [0, 1], keeps the peaks whose intensity is greater than *K*% of the highest peak intensity in S.

All considered standard SSM scoring functions are calculated based on the matched peaks between two spectra. Given two spectra, *S* = {(*MZ*_*i*_, *I*_*i*_)}_*N*_ and *S*′ = {(*MZ*′_*j*_, *I*′_*j*_)}, we need to obtain a pairing *PM* between a subset of peaks in *S* and a subset of peaks in *S*′ for SSM functions. The pairing *PM* is a set of peak pairs where the following conditions hold: (1) 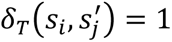 for any peak pair 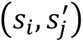 ∈ *PM*, and (2) each peak in either spectrum is used at most once in *PM*. We obtain the paring *PM* by maximizing the following function:

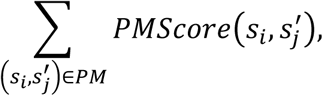

where *PMScore*(·) is a scoring function for matched peaks and depends on SSM functions, as defined below. A greedy method was used to find *PM* from 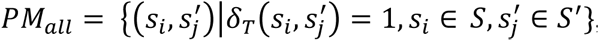, the set of all peak matches between *S* and *S*^′^. Note that *PM* ⊆ *PM*_*all*_, because a peak can match more than one peak from another spectrum (details are provided in Supplementary Algorithm S1.

##### Supplementary Algorithm S1.

A greedy method for finding the pairing *PM* between two spectra

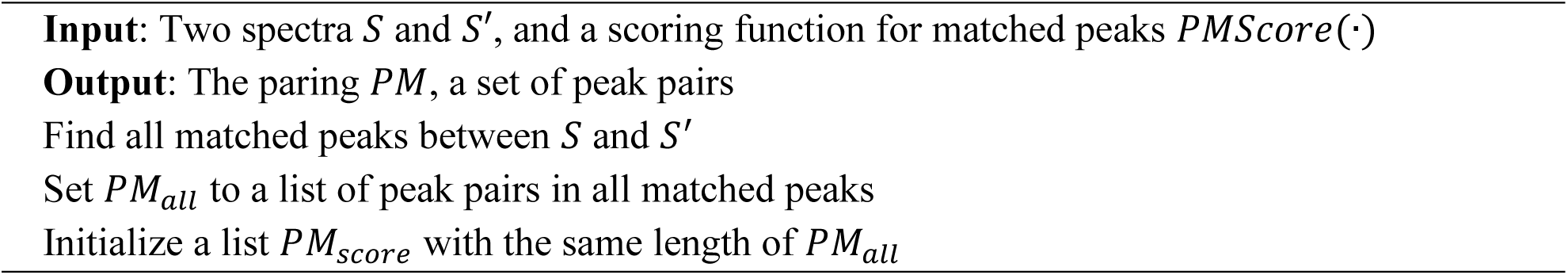
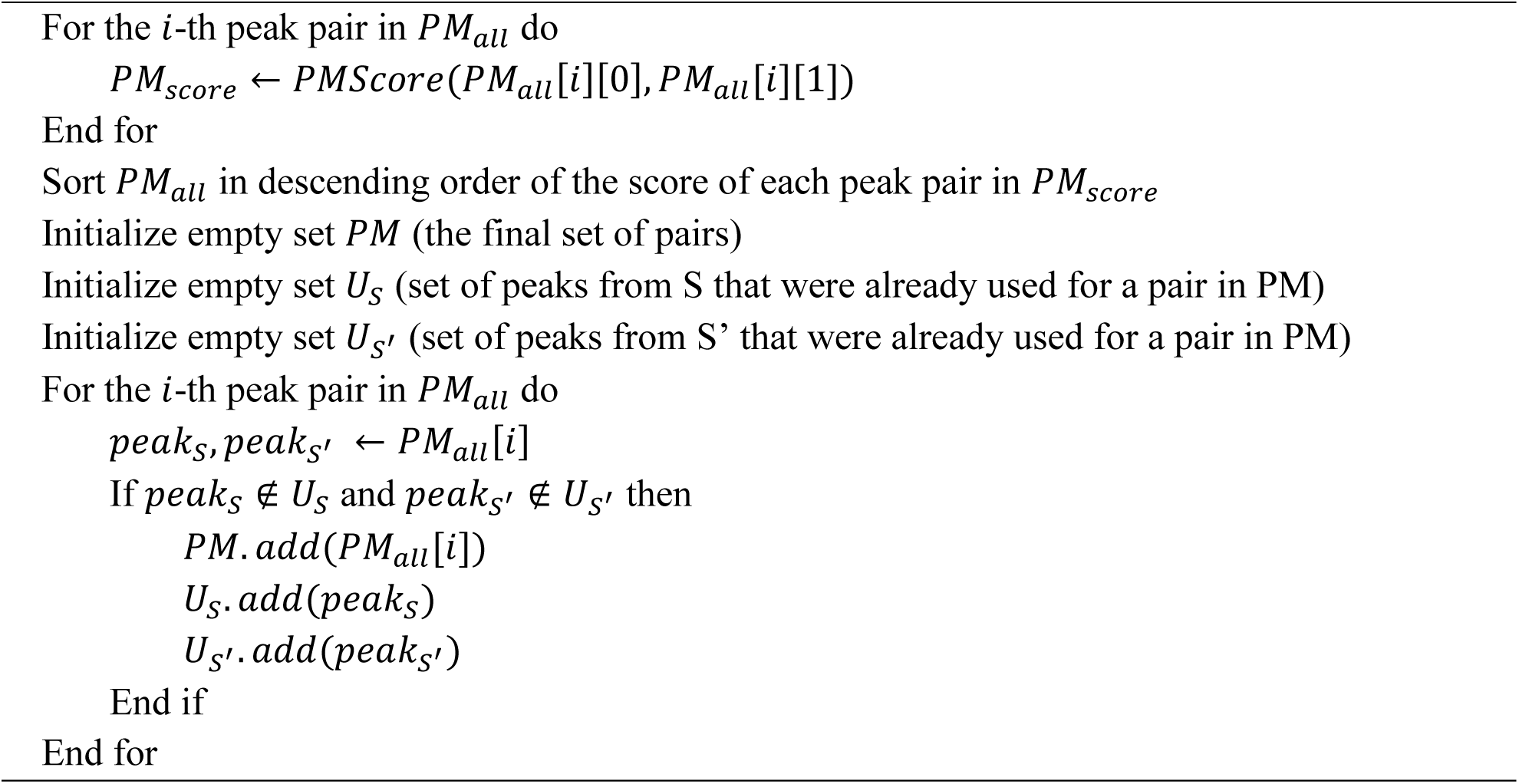

The most common SSM function is the normalized dot product, which is equivalent to *cosine similarity* when the vectors are pre-scaled to Euclidian norm 1. The cosine similarity between two spectra is defined here as

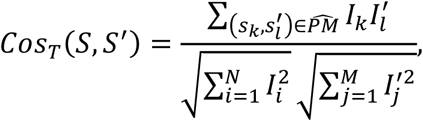

where 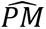 is the paring given by Supplementary Algorithm S1 for *S* and *S*^′^, with 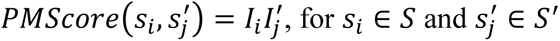.

Projected cosine^1^ computes the cosine similarity between *S*′ and a subset of *S*, from which all peaks not found in *S*′ are excluded from the score calculation (by re-normalizing S using only the intensities of its matched peaks).

The third SSM function is the symmetric form of relative entropy known as Jensen-Shannon divergence (JS-divergence). It considers a spectrum *S* = {(*MZ*_*i*_, *I*_*i*_)}_*N*_ as a distribution of mass per charge (*mz*) values of the peaks in *S*, where the probability of a peak occurring at *MZ*_*i*_ is

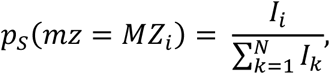

which is equivalent to the peak intensity after L1 normalization. The entropy of S is then calculated as follows:

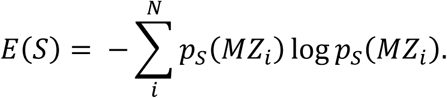

Consider two spectra, *S* = {(*MZ*_*i*_, *I*_*i*_)}_*N*_ and *S*′ = {(*MZ*′_*i*_, *I*′_*i*_)}_*N*_ with peak intensities L1-normalized such that 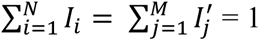. First, these two spectra are combined to create a new spectrum 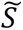, based on the paring 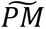. 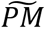 is obtained by Algorithm 2 with 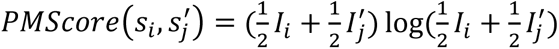 for *s*_*i*_ ∈ *S* and 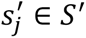. 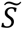 is the union of (1) 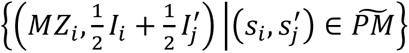 and (2) the rest of the peaks in both spectra that are not in 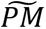. Then, the JS-divergence similarity score is defined as

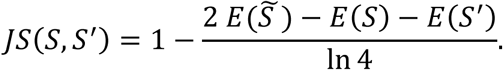

#### Supplementary Text S2: Differences between SpectraST/EntropyScore and standard SLS scoring functions

Compared to the benchmark pipeline, the spectrum matching scoring function in current SLS methods can incorporate more complex preprocessing steps and can have postprocessing steps to re-normalize spectrum matching scores with more information than just two spectra.

For SpectraST^2^, in the preprocessing step, query and library spectra are filtered with both *WF*_5,50_and *TOP*_50_, and the peak intensity of each peak is then replaced by 51 minus the peak rank. It then employs cosine similarity as an SSM function. In the postprocessing step, cosine scores for each query spectrum are transformed into probability scores based on the mean and standard deviation of the square-rooted cosine scores from the nontop-scoring library-matched SSMs for each query spectrum after excluding those with the five highest and lowest cosine scores.

EntropyScore^3^ employs JS-divergence for spectrum matching and introduces an adaptive preprocessing step that non-linearly transforms the peak intensities for both query and library spectra, according to the spectra’s entropy before using the JS-divergence scoring function. For a given spectrum *S* = {(*MZ*_*i*_, *I*_*i*_)}_*N*_, the intensity of the *i*th peak of the preprocessed spectrum 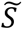 is

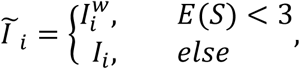

where *w* = 0.25(1 + *E*(*S*)).

##### Supplementary Algorithm S2.

Balancing candidate positives and guaranteed negatives

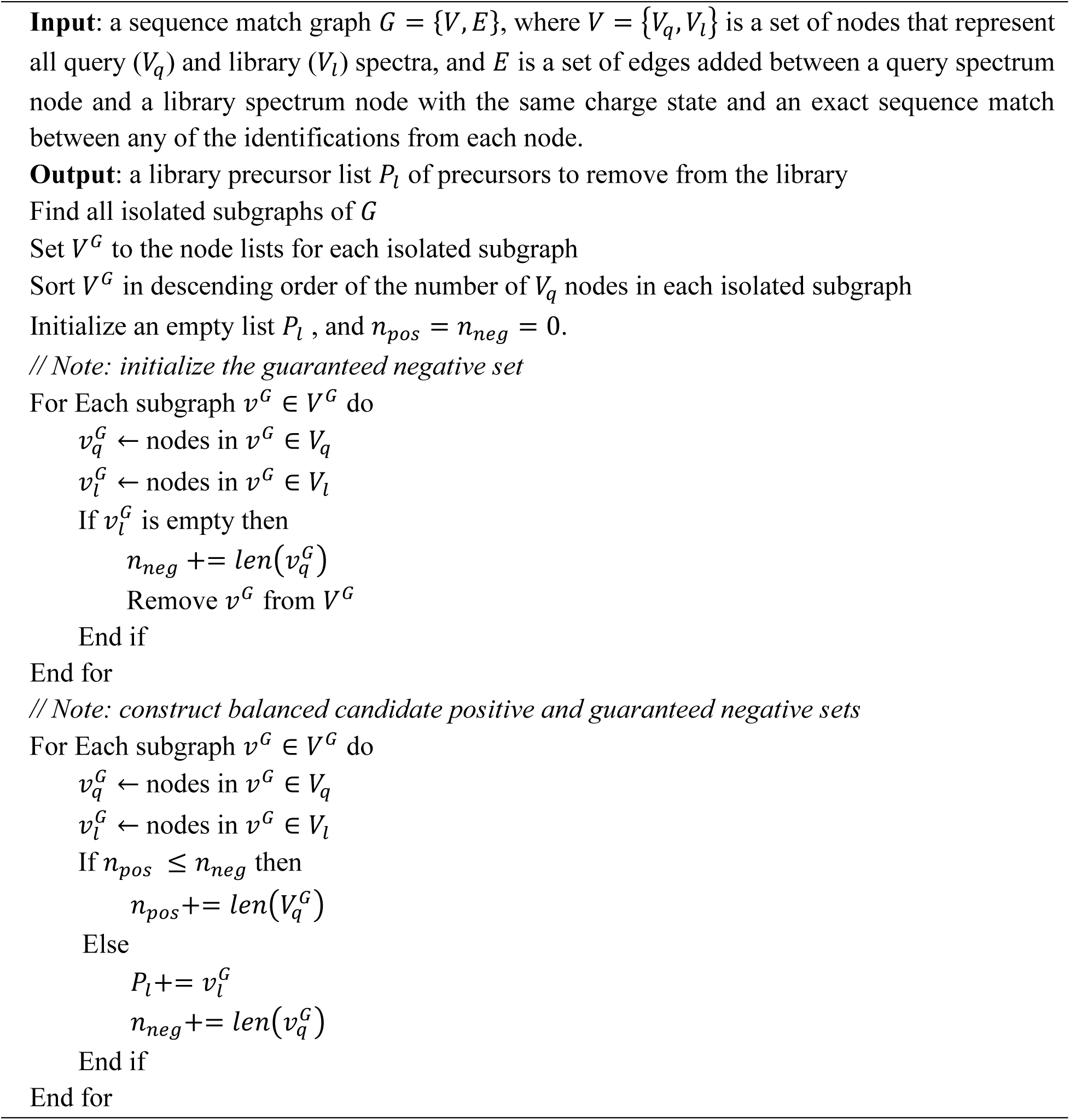

---

**Supplementary Figure S1.**
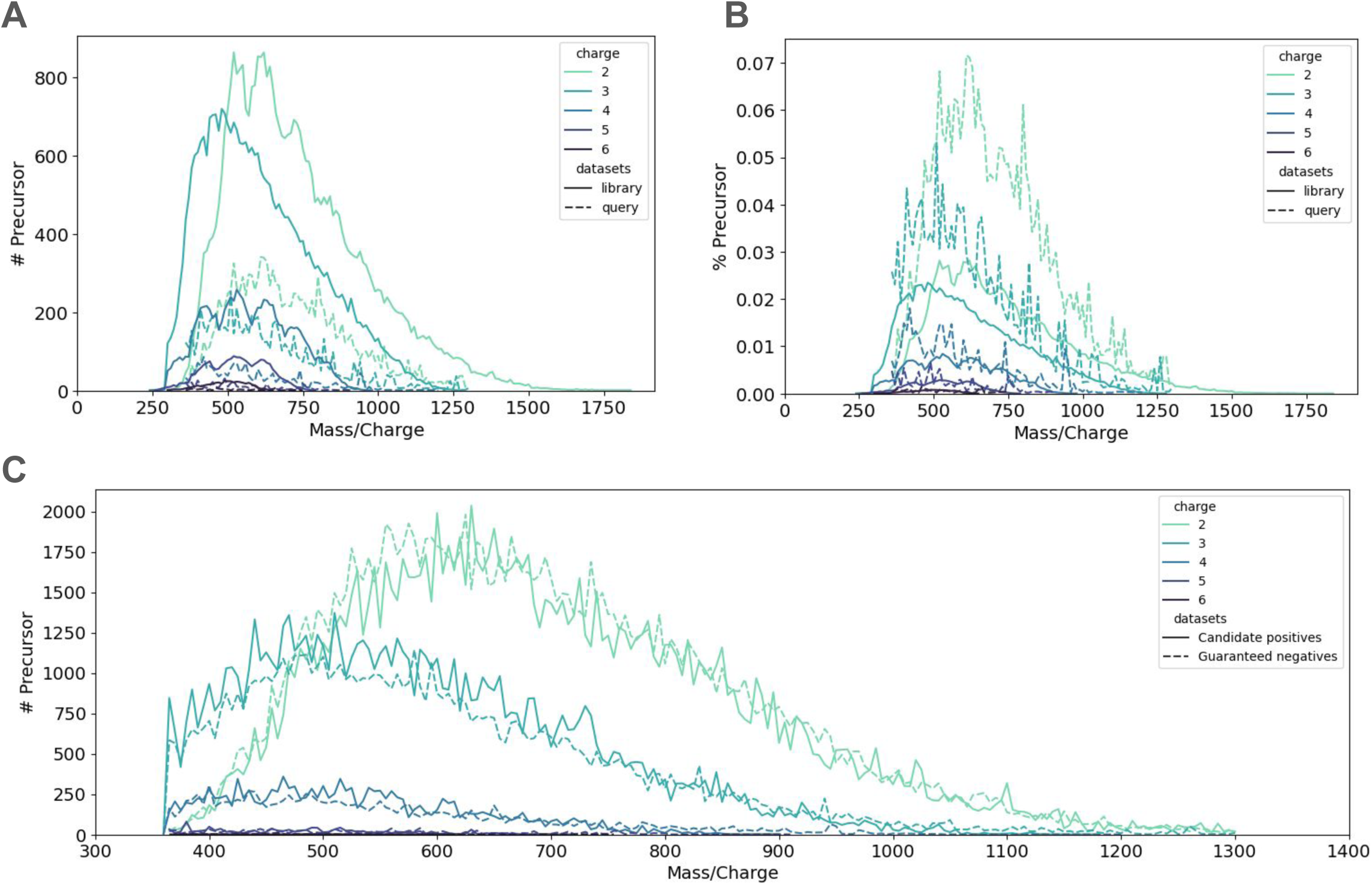
Precursor Mass/Charge distribution per charge state. (A) Number of precursors and (B) Percentage of precursors in query sets and spectral libraries, using bin size of 0.05 Da. Note that the figures only show the values at every 10 Da for better visualization. (C) Number of query precursors in the candidate positive and guaranteed negative sets with the bin size of 5 Da.

**Supplementary Figure S2.**
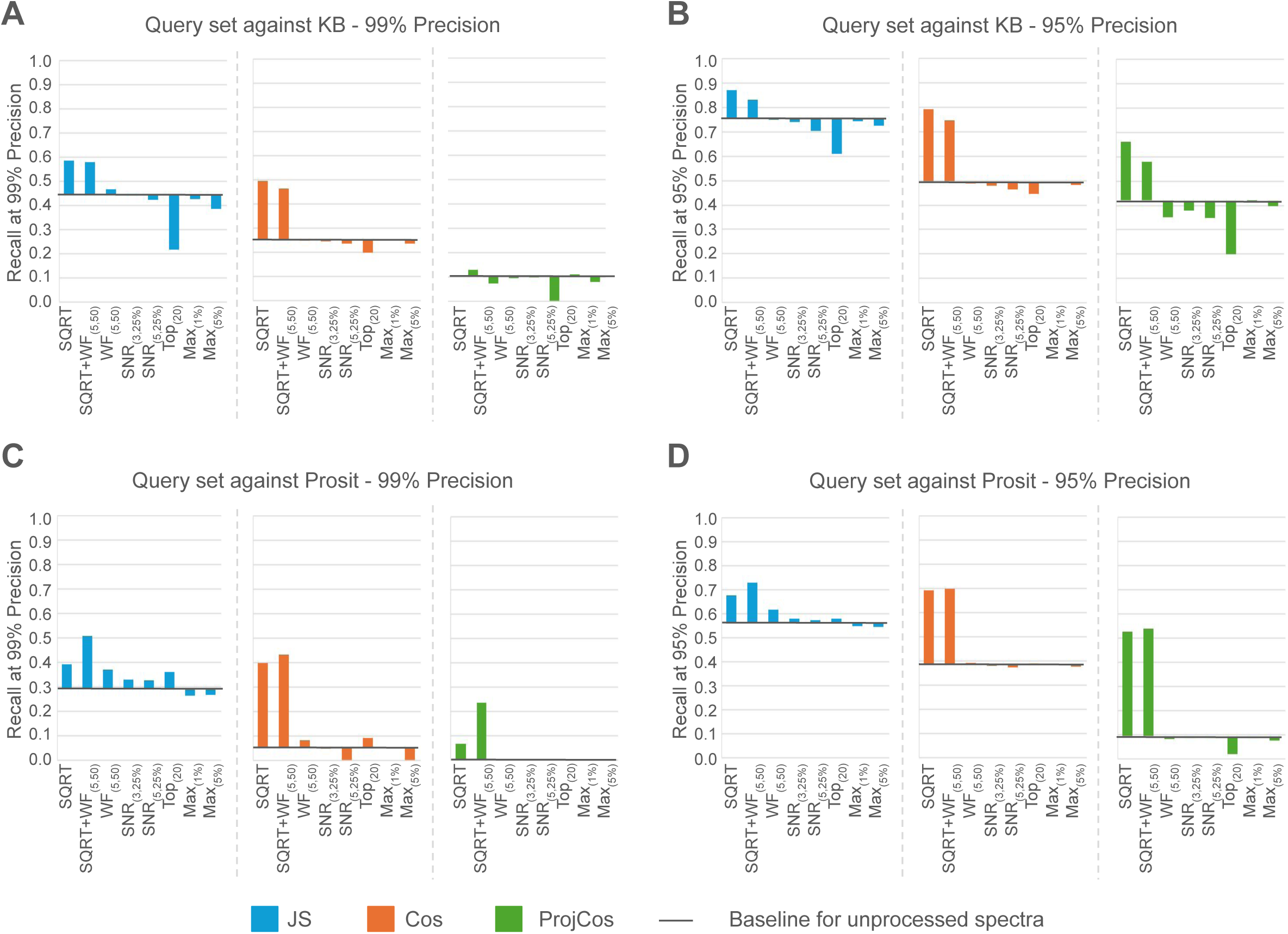
Additional results on the impact of preprocessing methods. The changes in recall after applying different preprocessing methods or their combinations for JS-divergence (blue), Cosine (orange) and Projected-cosine (green) searching query set against (A/B) the KB library and (C/D) the Prosit library at 99%/95% precision.

**Supplementary Figure S3.**
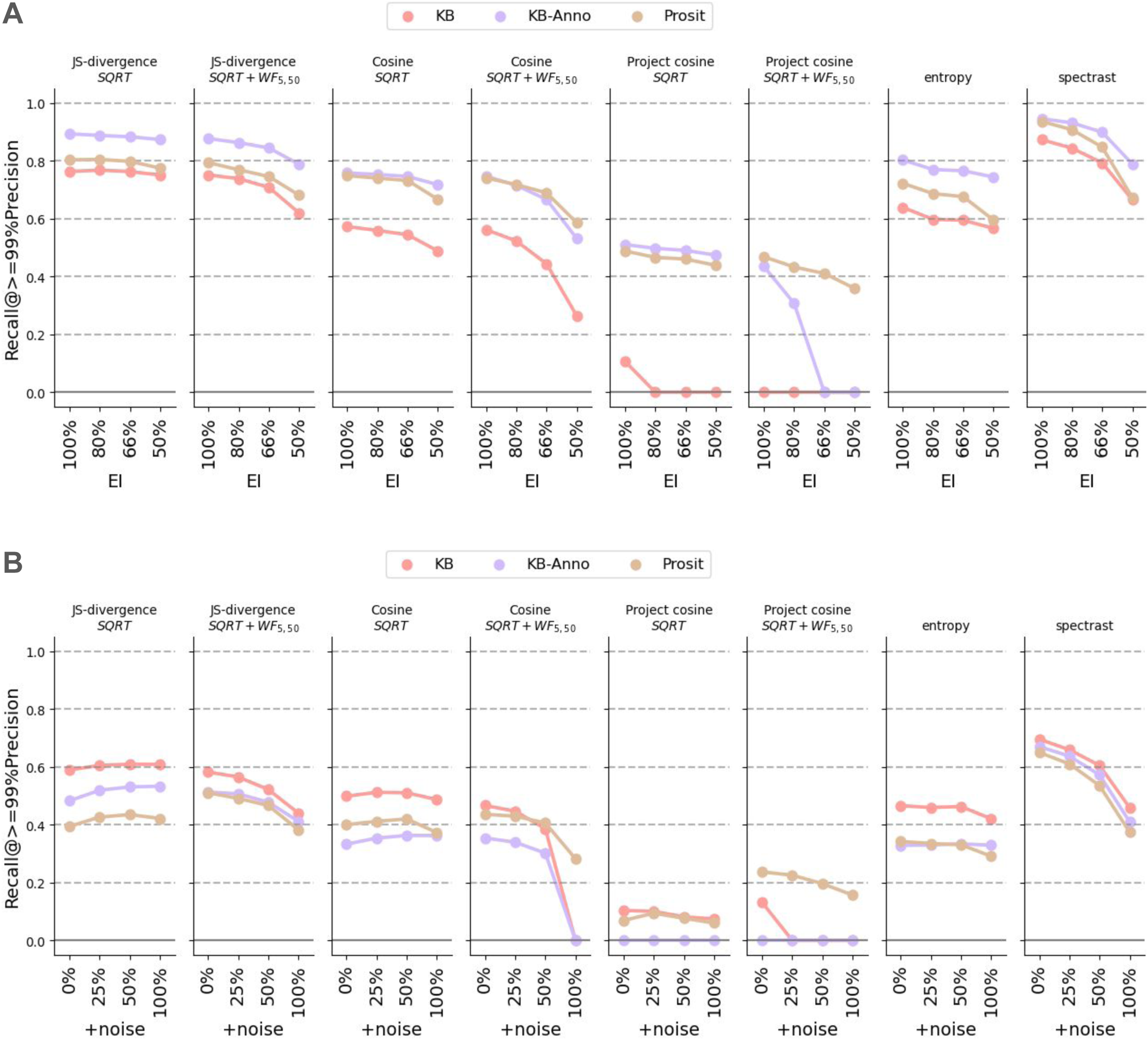
Additional results on the Impact of noise. Recall at 99% precision when searching (A) clean query spectra and (B) experimental query spectra with additional noise added at 0%, 25%, 50% and 100%. Note that when adding X% noise, the percentage of explained intensity is 100/(100+X).

**Supplementary Figure S4.**
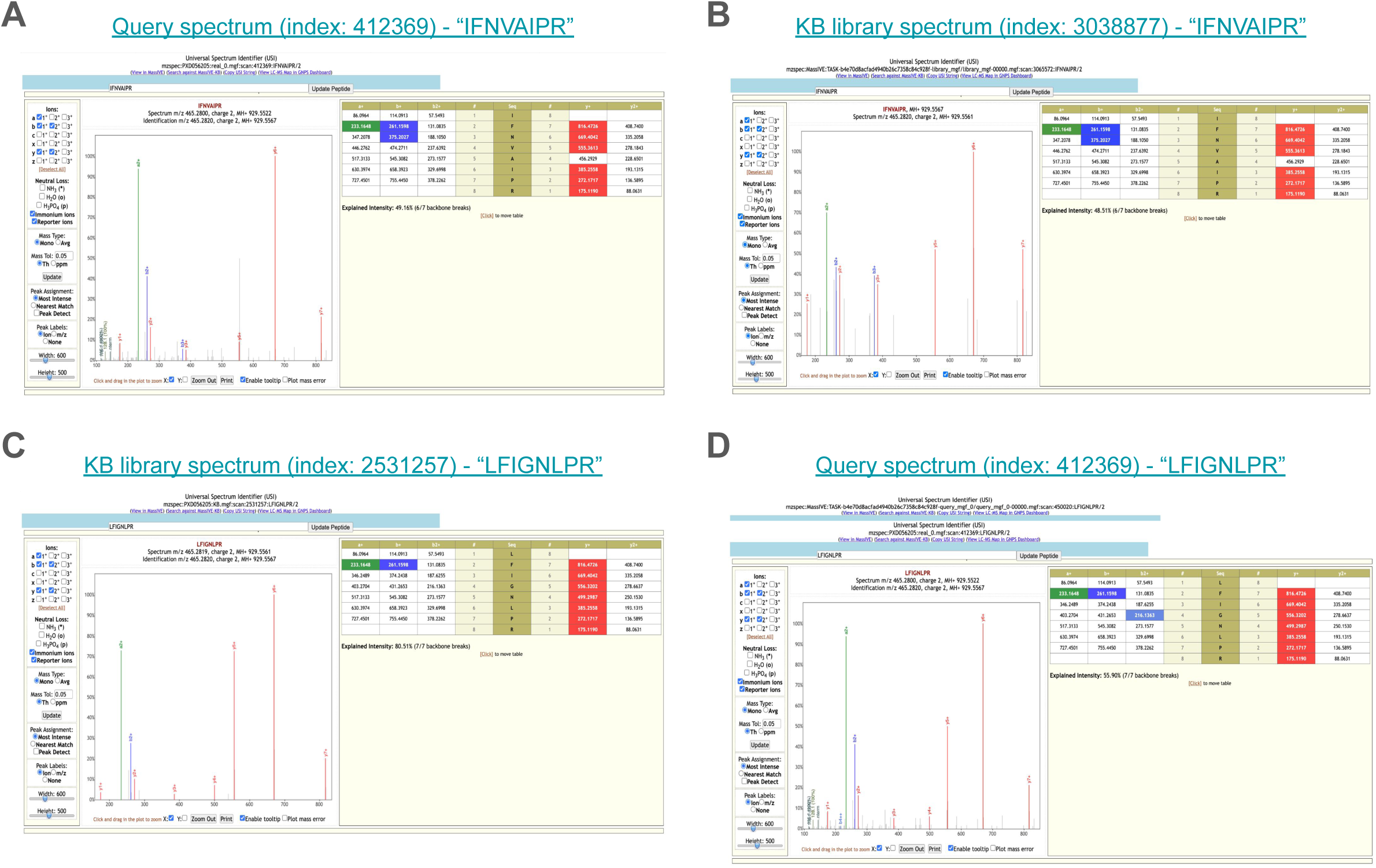
An example of the query spectrum from multiple peptides, which causes different methods to select different top matches. (A) The query spectrum, which an index of 450020, is identified as “IFNVAIPR” by MSGF+. When searching this query spectrum against the experimental library KB, the top-scoring library spectra given by (B) JS-divergence and SpectraST is identified as “IFNVAIPR” with <u>23 matching peaks</u> and (C) Cosine and the Entropy score is identified as “LFIGNLPR” with <u>15 matching peaks</u>, at 99% precision. (D) It seems that the spectrum in Figure S-4A might also come from peptide “LFIGNLPR”.

**Figure S5.**
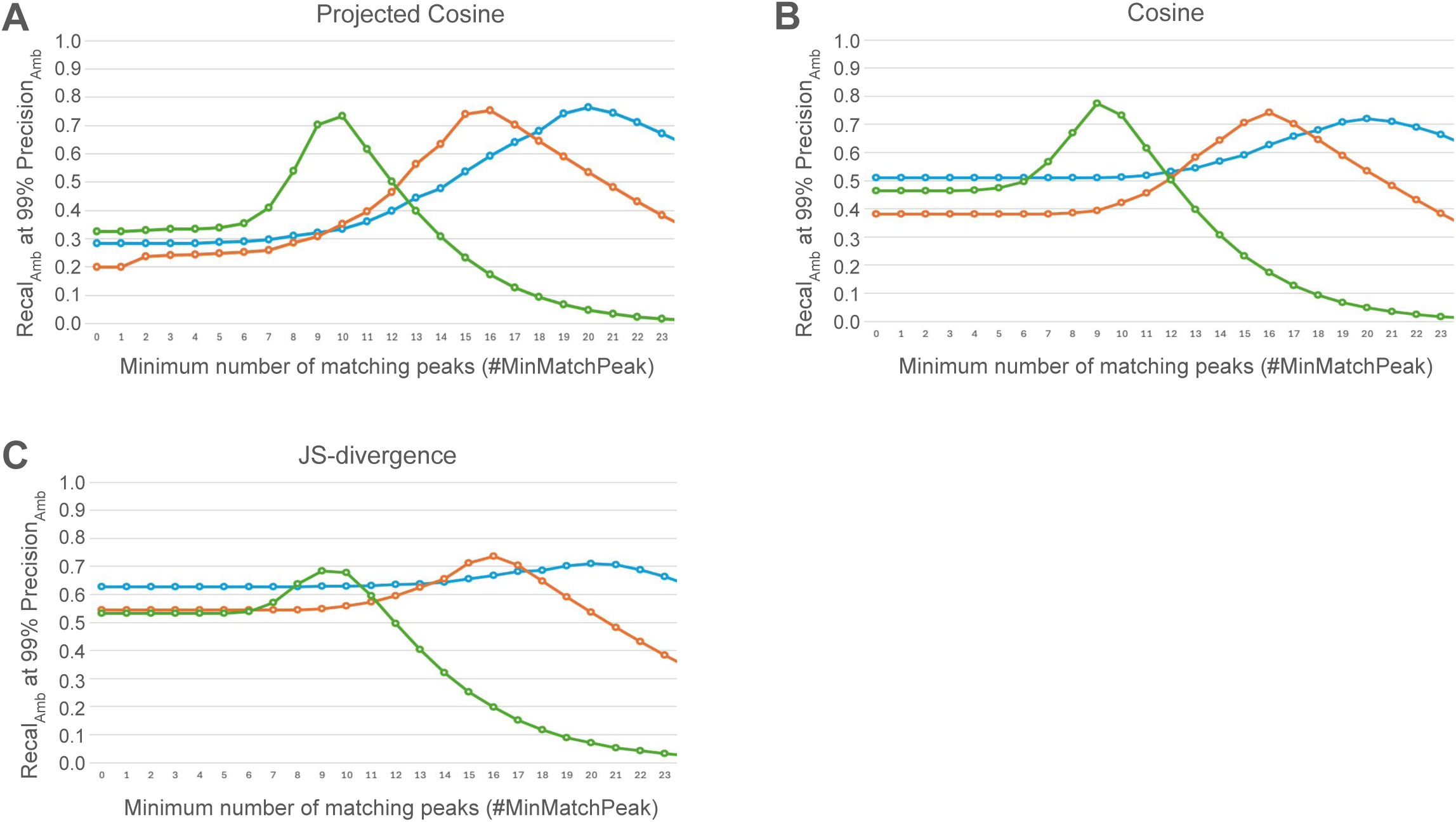
Impact of the minimum number of matching peaks. The ambiguous recall at 99% ambiguous precision for (A) Projected Cosine, (B) Cosine and (C) JS-divergence using the Prosit (green), KB-Anno (orange) and KB (blue) libraries. All query spectra are processed with the square root transformation (SQRT) and the window filter (WF_5,50_), and library spectra only with SQRT.

